# Beyond human-likeness: Socialness is more influential when attributing mental states to robots

**DOI:** 10.1101/2023.10.05.560273

**Authors:** Laura E. Jastrzab, Bishakha Chaudhury, Sarah A. Ashley, Kami Koldewyn, Emily S. Cross

**Affiliations:** Institute for Cognitive Neuroscience, School of Human and Behavioural Science, Bangor University, Wales, UK; Institute for Neuroscience and Psychology, School of Psychology, University of Glasgow, UK; Division of Psychiatry, Institute of Mental Health, University College London, UK; Chair for Social Brain Sciences, Department of Humanities, Social and Political Sciences, ETHZ, Zürich, Switzerland

**Keywords:** Social robotics, Second-person neuroscience, Social Cognition, & Mentalizing

## Abstract

We sought to replicate and expand previous work showing that the more human-like a robot appears, the more willing people are to attribute mind-like capabilities and socially engage with it. Forty-two participants played games against a human, a humanoid robot, a mechanoid robot, and a computer algorithm while undergoing functional neuroimaging. Replicating previous studies, we confirmed that the more human-like the agent, the more participants attributed a mind to them. However, exploratory analyses revealed that beyond humanness, the perceived *socialness* of an agent appeared to be as important, if not more so, for mind attribution. Our findings suggest that top-down knowledge cues are just as important, if not more so, than bottom-up stimulus cues when exploring mind attribution in non-human agents. While further work is now required to test this hypothesis directly, these preliminary findings hold important implications for robotic design and to understand and test the flexibility of human social cognition when people engage with artificial agents.

## Introduction

Robots have sparked curiosity and been romanticised in popular culture since von Kempelen’s “Chess Turk” was introduced in 1769. In the mid-20^th^ century, Alan Turing formalised the philosophical debate as to whether “machines think”,^1^ a question that continues to captivate many philosophical and science fiction writers. With the present study, however, we ask what might be thought of as the *opposite* question: namely, regardless of whether robots think, do *we humans* perceive robots as having minds of their own? If so, do we do so primarily based on how human-like the robot looks, or does its perceived socialness also matter?

Robots are already commonplace in assembly lines, factories, and dangerous jobs such as pipeline and fuel tank inspections, as well as underwater and space exploration.^2,3^ As the deployment of robots in these contexts grows, so does their introduction to social and leisure domains, aiding people with, for example, surgeries in healthcare, serving customers in restaurants, learning in schools, and supporting adults who need help with daily living skills (for example, ^4–8)^. Robots’ roles in our day-to-day lives so far, however, are typically “single-use” (e.g., robot vacuum cleaners or a robot check-in assistant at a hotel), and the ability of even the most sophisticated social robots to engage us socially is still far removed from depictions in science fiction novels and films.^9,10^ Rapid advances in hardware and artificial intelligence are expected over the coming decades, making this a crucial time to examine human engagement with robots. This is particularly true in the social domain if we are to develop machines that can indeed engage and collaborate with humans in complex social contexts.

As adults, humans typically and intuitively think of other humans as having a mind, thoughts, and intentions that are different from their own, a skill known as mentalizing.^11,12^ Mentalizing is important for social interactions, allowing us to read and react to others’ unspoken mental and emotional states, and their intended actions.^11^ Neuroimaging studies have used implicit (e.g., economic games) and explicit (e.g., mind-in-the-eyes) tasks to probe human brain activity associated with mentalizing (for a review, see ^13^). This work has identified the so-called mentalizing network, a group of brain regions thought to support thinking about others’ minds. The core regions reliably included as part of the mentalizing network include bilateral temporal-parietal junction (TPJ), medial prefrontal cortex (mPFC), and Precuneus (PreC) but engagement of additional brain regions, including posterior superior temporal sulcus (pSTS), temporal poles, and posterior cingulate cortex (PCC), have also been implicated.^13–18^

The mentalizing network is readily engaged during interactions with other humans, especially when trying to predict their future actions. Very few neuroimaging studies, however, have directly addressed the extent to which mentalizing brain regions, which have ostensibly evolved to interpret other people’s actions and intentions, also process non-human social partners such as robots. Understanding whether humans mentalize about robots is important for at least two reasons. First, the more we attribute a mind to robots, the more likely we are to interact with and engage with them socially.^19–21^ Second, examining mentalizing in response to robot social partners tests the flexibility of our social cognitive system by assessing the extent to which a system that evolved to support interactions with fellow humans can be engaged during interactions with non-human agents (in this case, robots).^22^ Prior neuroimaging studies studying the extent to which humans mentalize about robots have used empathy tasks,^23,24^ spatial cueing tasks,^21,25^ and economic games.^26–28^ Several of these studies demonstrate that human—robot interactions (HRI) activate the mentalizing network, but to a lesser degree than human—human interactions (HHI).^26,27,29^

One influential theory that might help to explain the pattern of activity reported so far is the ‘like-me’ hypothesis,^30^ which posits that the more human-like a non-human agent appears, the more readily social brain networks are engaged. Indeed, behavioural data generally support this idea. For example, the more human-like a robot appears, the more a human user will expect that robot to behave like a human.^31^ Furthermore, a robot’s appearance influences our assumptions about its behavioural capabilities^32–34^ and the extent to which we attribute intentionality or a mind to them.^19–21,35,36^ Likewise, the degree to which we anthropomorphize robots (or attribute human-like qualities to them) has also been found to depend upon a robot’s human-like appearance and behaviour.^37–40^ Given the behavioural evidence, it is perhaps not surprising that similar results are found when examining socio-cognitive brain systems. For example, Krach and colleagues^27^ reported that the increasing human-likeness of game partners’ physical features was associated with increasing engagement of mentalizing network regions during an implicit mentalizing task (in this case, an iterative prisoner’s dilemma game). Together, behavioural and brain imaging findings support the idea that the human-likeness of an interactive partner’s appearance plays a key role in engaging socio-cognitive processes like mentalizing. However, emerging evidence raises the possibility that human-likeness alone may not fully explain which robots are seen as more desirable social partners and, thus, which robot features might be most effective at eliciting the strongest human-like social-cognition processes.^31,41,42^ The influence of a robot’s *social* features, per se, on human perception and engagement is an emerging area of research that will benefit from expertise from the Human Robot Interaction (HRI), social robotics, and cognitive neuroscience communities.

In the current study, we sought to replicate prior findings that the mentalizing network increases in responsiveness as the appearance of robots increases in human-likeness. In additional exploratory analyses, we sought to explore the extent to which a partner’s perceived *socialness* (independent from human-like physical features) might also contribute to this process. To do so, we used an established implicit mentalizing task where participants play rock-paper-scissors (RPS)^43^ against a human and several artificial agents. We followed an experimental design like that reported by Chaminade and colleagues.^26^ The RPS game itself is familiar across cultures and age groups, and if it is unfamiliar, it is easy to learn. Also like Chaminade and colleagues,^26^ we used videos of game partners to increase the sense of live interactions during game play. We controlled wins and losses across all game partners, and explicitly told participants that the robot competitors had been endowed with artificial intelligence and would play strategically. Similar to Krach and colleagues,^27^ we included two robotic partners that differed in their human-like appearance. One robot appeared humanoid, with clear human-like features including a body, torso, arms, hands, fingers, head and eyes. The other was a mechanoid robot, which had expressive eyes but no other human-like physical features (see Fig 5). Importantly, both the humanoid and mechanoid robots in our study are designed to engage people with socially interactive behaviours.

From prior data, we expected that both robots would engage the mentalizing network, though to a lesser extent than the human game-partner. Indeed, we preregistered a prediction that the magnitude of response of core brain regions within the mentalizing network (specifically TPJ, mPFC, and Precuneus) would linearly increase as game partners increased in human-like appearance. We further explored the extent to which participants found each robotic game partner fun, sympathetic, competitive, successful, strategic, intelligent, and competitive. Here we hypothesized, again based on previous findings^26,27^ that these factors would increase with increasing human-likeness. Finally, in an exploratory analysis, to address questions related to participants’ perceptions of the socialness of the different game partners, we reversed the order of the robots in our linear contrast models, allowing us to test the extent to which this “perceived socialness’’ might explain differences in the engagement of the mentalizing network across game partners *better* than simply the agents’ physical appearance.

## Results

### Neuroimaging Results

#### Socialness and human-likeness influence mentalizing but socialness is more robust

##### Pre-registered

Repeated-measures ANOVAs with game partner as a within-subjects factor was significant in several key mentalizing ROIs during game play (bilateral TPJ and left middle frontal gyrus (lmFG)), as well as, bilateral pSTS. Follow-up paired sample t-tests in bilateral pSTS and lTPJ revealed that this was largely driven by higher activity in response to the human compared to all other conditions, suggesting that these regions are more reliably engaged by human than artificial stimuli (see Supplementary Table 5). Right TPJ was an exception, in that, while the human significantly differed from both robots, no significant difference between the human and computer was found. No other significant comparisons during gameplay and within these ROIs remained after correcting for multiple comparisons.

Results from the pSTS revealed significant differences between game players while playing the game (rpSTS: F(3, 123) = 12.39, p < 0.001, np = 0.23; lpSTS: F(3, 123) = 6.96, p < 0.001, np = 0.15), which was unexpected as there were no visual differences during game play across the 4 conditions.

Contrary to our expectations, mentalizing regions were not activated above baseline during the RPS games. Average activity across the group was close to zero or, indeed, slightly negative across nearly all conditions (see Figs 1, S2 & Table S5).

**Fig 1.**
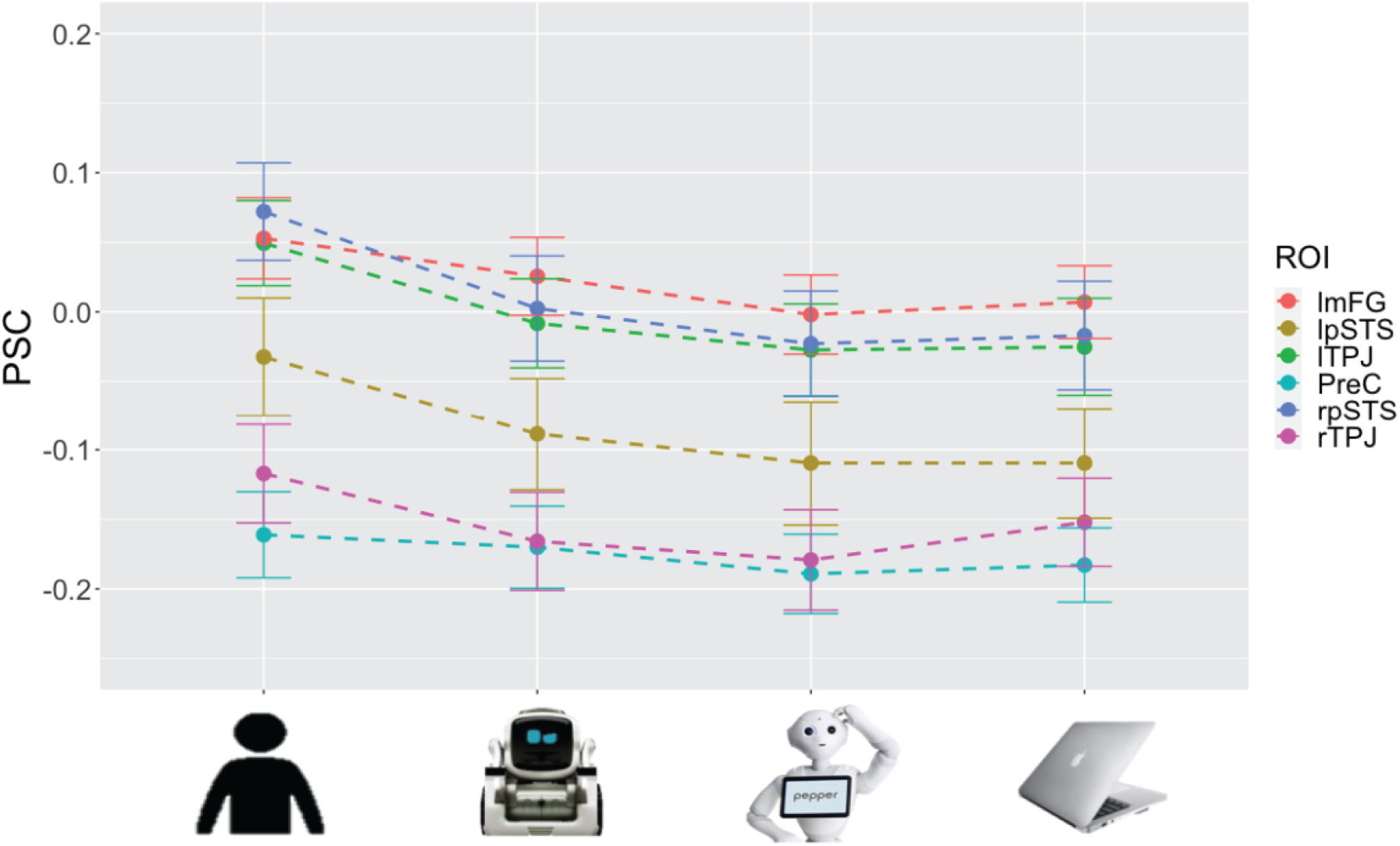
Average percent signal change (PSC) during gameplay in mentalizing ROIs and pSTS with significant within subject rmANOVA (Error bars are SEM).

##### Exploratory

Additionally, the pSTS revealed strong significant differences across game partners while participants watched the introductory video (video 1) of each game partner before playing commencing each game series (rpSTS: F(3, 123) = 29.40, p < 0.001, np = 0.42; lpSTS: F(3, 123) = 13.26, p < 0.001, np = 0.24). While none of the other ROIs revealed significant pairwise differences between either robot and the computer, there was a significant difference between MR and CP in rpSTS (and approached significance in lpSTS) during the video preceding gameplay (rpSTS: p < 0.001, d = -0.73; lpSTS: p = 0.056, d = -0.32; See Supplementary Table S5).

#### Linear effect of human-likeness in mentalizing ROIs during gameplay

##### Pre-Registered

All mentalizing ROIs which revealed a significant within subject effect of partner (Bilateral TPJ, lmFG, and bilateral pSTS) also revealed a significant linear within-subjects contrast effect of human-likeness (HP > HR > MR > CP), as predicted (see Table S5).

##### Exploratory

We explored whether changing the order of the robots in the within-subject contrasts according to socialness ratings further bolstered the linear effect (HP > MR > HR > CP, see Table S5). Results from behavioural ratings suggested that socialness (as assessed by perceived fun, competitiveness, and sympathy, see below) models were improved by reversing the order of the robots. Indeed, across ROIs, the mechanoid robot evoked numerically higher, though often not significantly so, responses than the humanoid robot. Despite the lack of statistically significant differences between the robots in pairwise comparisons, the linear effect of ‘socialness’ resulted in a larger effect size than the ‘humanness’ model, suggesting socialness may be even more important than humanness in mind attribution toward robots, as measured by engagement of brain regions associated with mentalizing.

#### The mechanoid is more similar to the human than the humanoid or computer

##### Pre-Registered

No FWE (p < .05) or uncorrected (p < .001) clusters survived simple whole brain contrasts between the humanoid or mechanoid and the computer (see Supplementary Table S4). There were no significant clusters during the [Humanoid (HR) > Mechanoid (MR)] but the inverse contrast revealed a significant cluster (k = 313) in nucleus accumbens (MNI: -4 10 -10). The [Human Partner (HP) > Computer Partner (CP)] contrast resulted in significant mentalizing clusters in bilateral TPJ, mFG, mPFC, precuneus, rpSTS, IFG, nucleus accumbens, and cerebellum.

To assess whether regions outside our pre-selected ROIs might be sensitive to Human-likeness, we tested whether any brain regions showed a pattern of activity such that Human Partner (HP) > Humanoid Robot (HR) > Mechanoid Robot (MR) > Computer Partner (CP). This analysis revealed that rTPJ, precuneus, mPFC, bilateral mFG, and nucleus accumbens all survived the FWE-corrected peak-level threshold.

##### Exploratory

When the human was compared to the humanoid and mechanoid robots, several regions associated with mentalizing were significant at the cluster level after FWE correction (see Fig. 2). The [HP > HR] contrast resulted in significant clusters in bilateral TPJ, precuneus, rmFG, rIFG, rpSTS after FWE corrections. The [HP > MR] contrast yielded significant engagement of rTPJ, precuneus, rpSTS, and cerebellum after FWE corrections.

**Figure 2.**
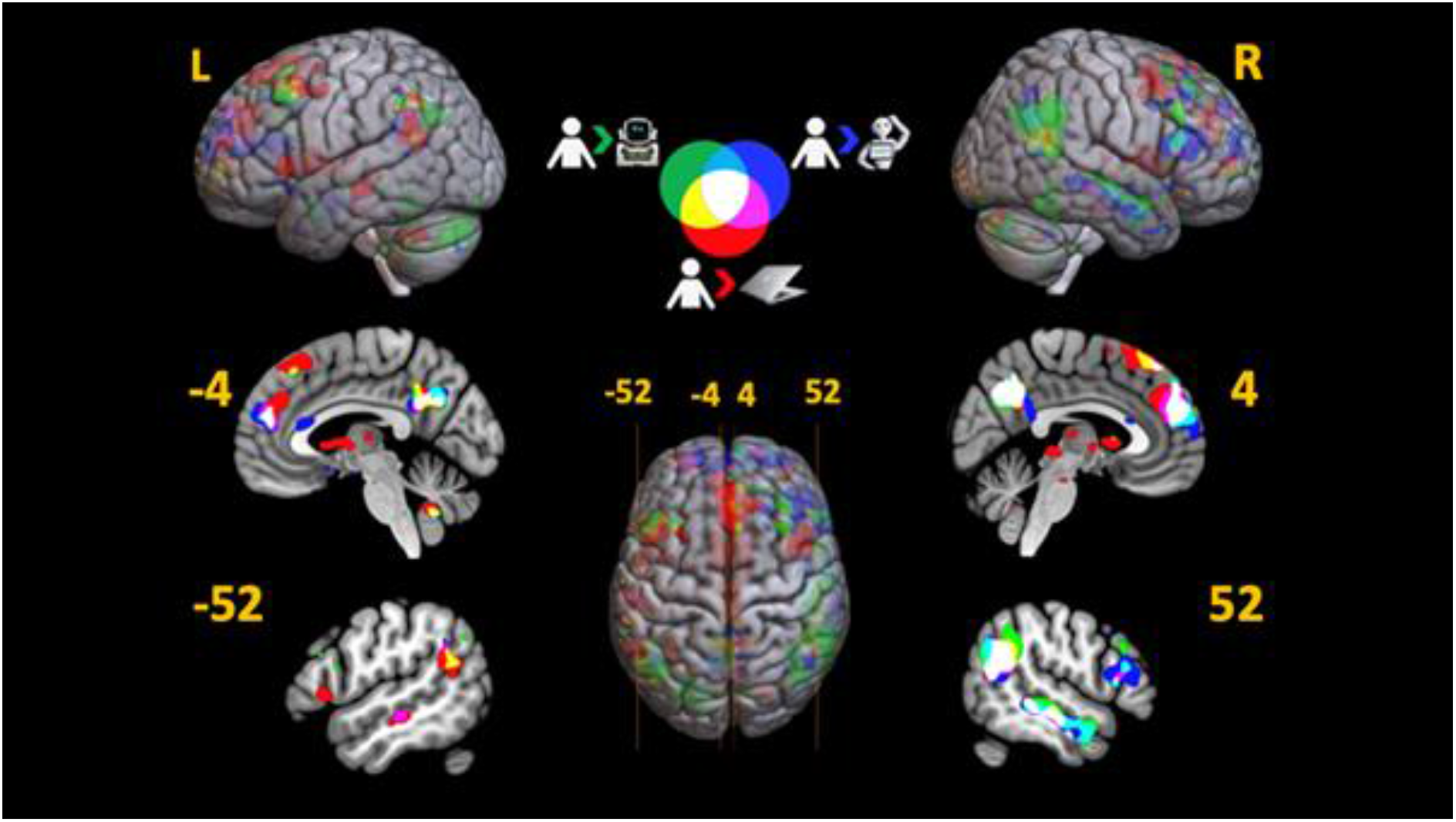
Whole brain T-map overlap analysis (Human > Computer (Red); Human > Humanoid (Blue); Human > Mechanoid (Green)). There were no significant whole brain differences between either robot and the computer condition nor between the robots themselves; the robots and the computer conditions may be perceived similarly. The Human-Computer contrast (red) revealed significant clusters in mentalizing regions (ie Precuneus, bilateral TPJ, and mPFC), similar with previous findings. There was less overlap between the Human-Mechanoid contrast (green) with the Human-Computer contrast suggesting that there may be differences in how participants attribute a mind to the mechanoid compared to the humanoid and computer.

In line with our socialness questions, we also tested whether any brain regions showed a pattern of activity if we reversed the order of the robots in our parametric analysis; i.e., so that the order was now: Human Partner (HP) > Mechanoid Robot (MR) > Humanoid Robot (HR) > Computer Partner (CP). Results revealed a similar pattern to both HP>CP and the HP>HR>MR>CP model above but now also included significant clusters in: bilateral pSTS, supplementary motor area, rIFG,& lTPJ. Please see Figure S1 and Table S4.

### Behavioural Results

#### Manipulation Check

During verbal debriefing with participants, six out of 42 neuroimaging participants questioned whether the videos were live during our verbal debriefing. Given this, we re-ran all behavioural and neuroimaging analyses with only the “true believers” (see OSF project page for details). Doing so did not change the findings in either degree or direction of significance. Therefore, the analyses are reported with the full sample, including the non-believers.

#### Debrief Questions: Mechanoid perceived as more social, but not intelligent, than the humanoid

##### Pre-Registered

All pairwise comparisons in this section were corrected for multiple comparisons (Bonferroni). Greenhouse-Geisser corrections were made if any rmANOVA was found to violate Mauchley’s tests of sphericity (see Figure 3, S3, & Table S6 for details from this section).

**Fig 3.**
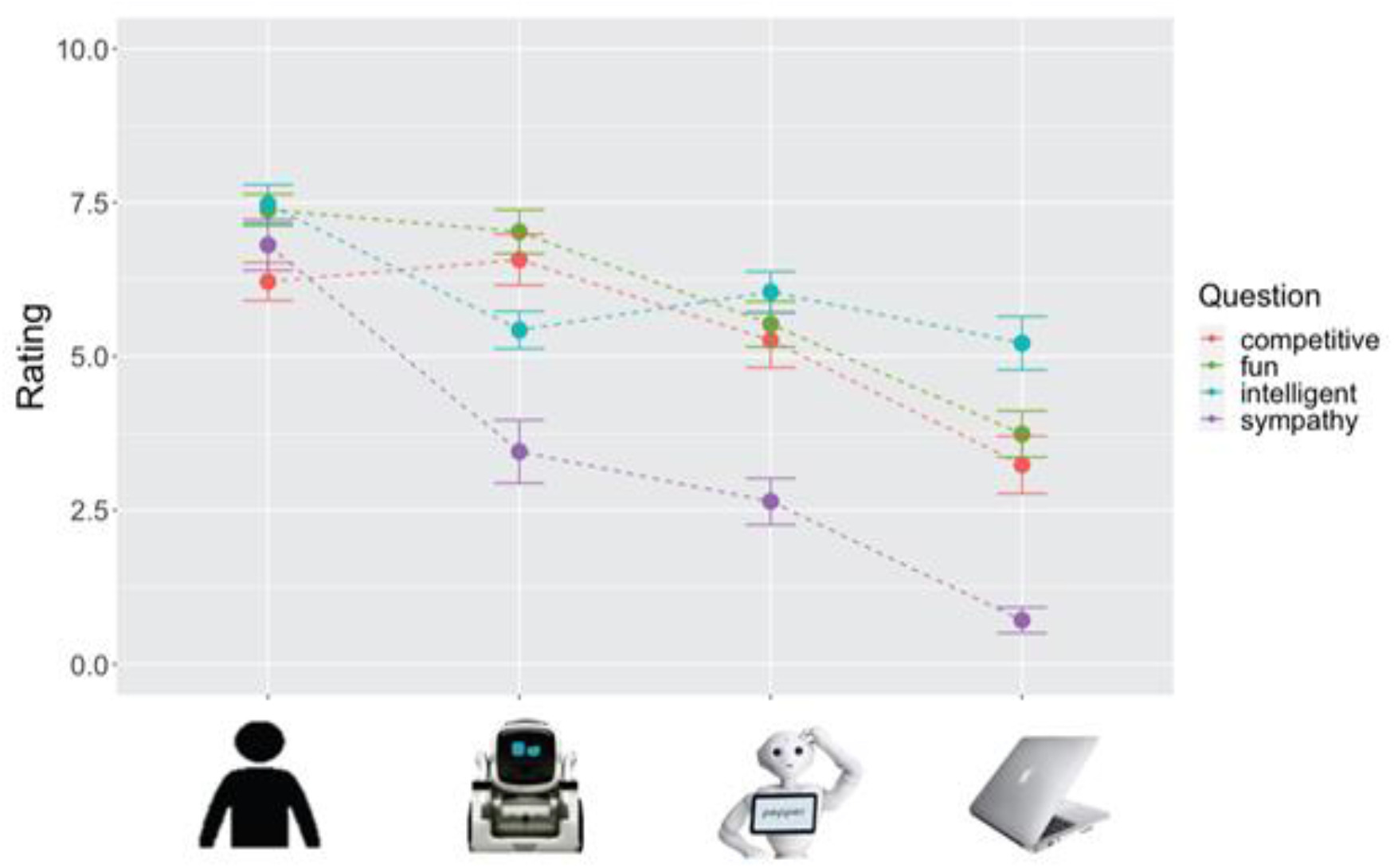
Average Likert (0-10) scale ratings of Debrief questions (Error bars are SEM).

We found no effect of perceived *success* in winning (F(3, 123) = 0.50, p = 0.685, ηp^2^ = .012) or *strategy* employed (F(3, 123) = 0.32, p = 0.811, ηp^2^ = .008) against each game partner, despite stressing to participants that the computer was using a random algorithm, while the other partners were all trying to win.

*Fun* (F(3, 123) = 33.90, p < 0.001, ηp^2^ = .453), *Competitiveness* (F(3, 123) = 17.24, p < 0.001, ηp^2^ = .296), *Sympathy* (F(2.50, 102.58) = 58.59, p < 0.001, ηp^2^ = .588; Greenhouse-Geisser corrected) and *Intelligence* (F(2.51, 102.91) = 12.16, p < 0.001, ηp^2^ = .229; Greenhouse-Geisser correction) were all significantly different amongst the four conditions and followed a significant linear pattern based on human-likeness.

##### Exploratory

However, *Fun*, *Competitiveness*, and *Sympathy*, revealed a stronger linear pattern based on socialness, wherein the two robots were reversed in the order of the within subject contrasts. However, only post-hoc tests on ratings of *Fun* and *Competitiveness* showed differences between robots, where mean ratings for the mechanoid robot were higher than for the humanoid robot (p=0.006 & p=0.049, respectively).

#### Inclusion of Others and Self (IOS): No difference in perception of closeness between the robots or a human stranger

IoS scores varied significantly between the 6 agents (F(3.70, 148.02) = 122.40, p < 0.001, ηp^2^ = 0.754). Pairwise comparisons of the computer, human game partner, and close friend significantly differed from all other agents and each other on the IoS, even after correcting for multiple comparisons (Bonferroni). Pairwise comparisons of the mechanoid robot, humanoid robot, and human stranger did not significantly differ from each other (see Fig. 4 & Table S7).

**Fig 4.**
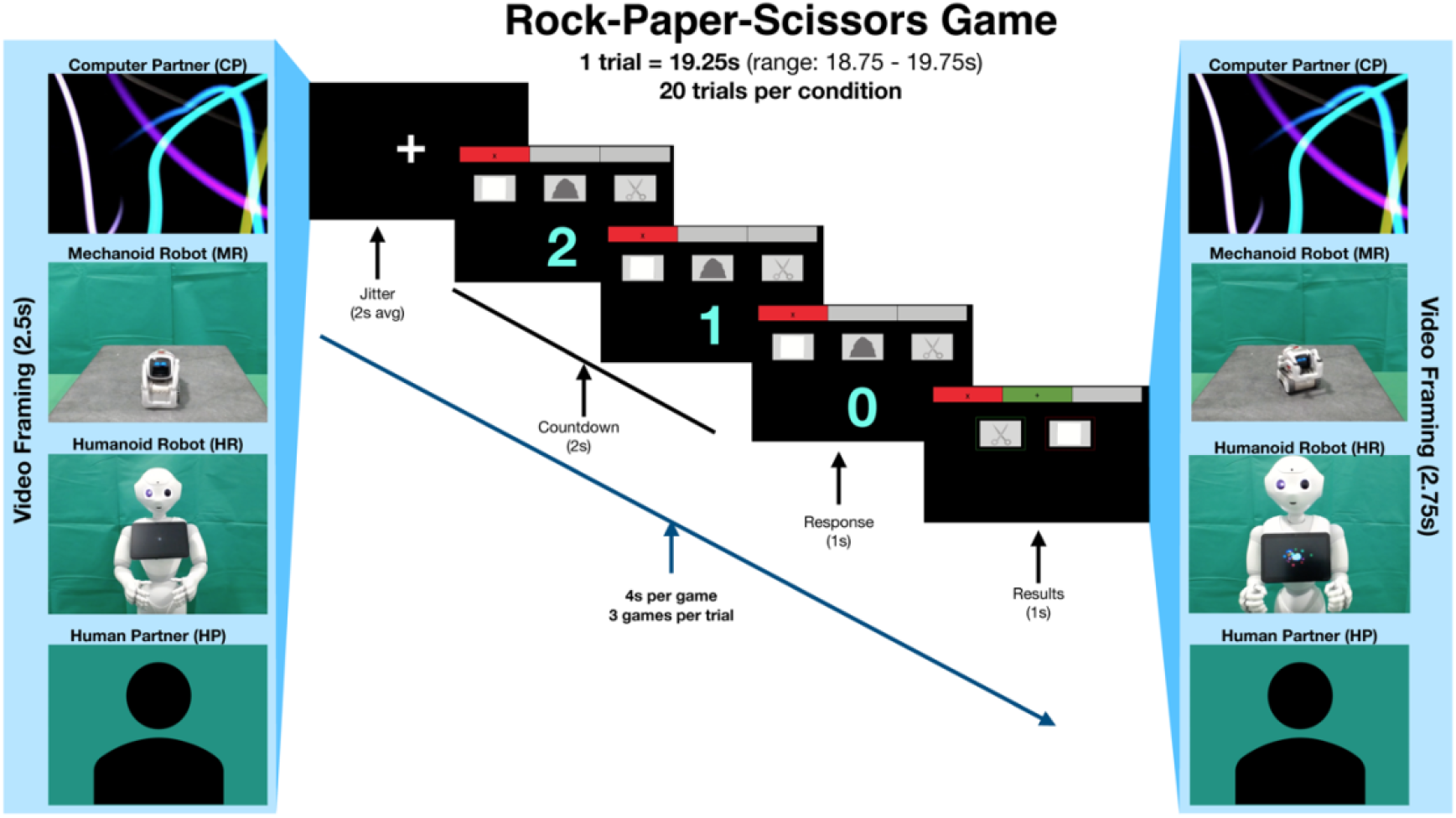
Experimental paradigm. Each game lasts 4 seconds and includes a countdown from 2 to 0; participants respond on ‘0’ and then the results of the game are shown to the participant before the next game in the series commences. Participants play 3 games against one opponent per trial. Each trial is preceded by a 2.5 second video of their opponent (video framing) and concluded by a 2.75 second (video feedback) of their opponent; each series of 3 games lasts 12 seconds without video framing and feedback (average of 19.25 second in total).

## DISCUSSION

With the present study, we have replicated and extended previous findings, demonstrating that both human likeness and perceived ‘socialness’ shape the extent to which participants engage mentalizing regions while playing games against robotic partners. We found that although human-likeness models showed increased theory-of-mind network engagement (as predicted and pre-registered), the socialness model was even more robust. While this analysis was exploratory and will require replication via hypothesis-confirming follow-up work, it is important for two reasons. First, it suggests that mentalizing processes during interactive exchanges (in this case, a game) are better predicted by how *social* we find our interaction partner, rather than being solely based on how human-like they look. This finding has the potential to update our models of how mentalizing systems can be engaged, particularly by non-human interactants. Secondly, the extent to which humans will ascribe mental states to robots is likely to become increasingly relevant as roboticists develop increasingly sophisticated embodied artificial agents designed to engage human users on a social level. Successful social interactions with such social robots will require people to think about how the robot “thinks”. A better understanding of the factors that influence mentalizing towards and about robots should lead to higher quality and more sustained long-term interactions with robots in social domains (e.g.,^42^).

As with the two previous neuroimaging studies on which we based our current study, we found increasing activation in mentalizing regions with increasing human-likeness.^26,27^ We also found similar behavioural ratings, showing that while participants did not perceive strategy and success differently across game partners (suggesting participants did not feel that they won or lost more against any one game partners), participants did perceive the game partners differently based on social factors like perceived intelligence, fun, competitiveness, and sympathy. However, unlike previous studies, we explored how these social factors might contribute to mind attribution and found that reversing the order of robots in our linear contrast models to reflect participants’ evaluations of socialness resulted in numerically stronger models than those based on the human-likeness of physical features alone.

### Quantifying and exploring human-likeness vs. socialness

While the human and social models were both significant and strong, one possibility for the numerically stronger social model is that the mechanoid robot was perceived as more social because it exhibited higher levels of hedonic factors (as rated by fun, competitiveness, and sympathy) than did the humanoid robot. This finding is consistent with participant qualitative perceptions and behavioural ratings of this same robot in recently published work.^44,45^ For example, in one scenario from our study, when the mechanoid robot lost the RPS series, it pouted and slammed its forklift on the table while moving around in circles in protest. Whereas, when the humanoid robot lost, it responded similarly to the human in a more measured manner, by lowering its arms and shaking its head and/or looking down in defeat. While these differences in personality and behaviour were not objectively measured in our study, others report that manipulating social features of robots such as personality,^46,47^ emotional arousal,^48^ and other hedonic features such as enjoyment and sociability^49^ can increase user engagement, acceptance, and/or satisfaction.

The neuroimaging evidence from this study supports both human likeness and socialness models when attributing mental states. Bilateral TPJ, bilateral pSTS, and lmFG showed significant increases with human-likeness and a numerically stronger linear increase with socialness. While we expected the whole mentalizing network and pSTS to show a similar response pattern, the exceptions were in mPFC, precuneus, and rmFG.

We were unable to clearly assess the role played by our mPFC, Precuneus, and rmFG ROIs in this study, as we found no significant differences to emerge between the agents during game play. However, a wealth of research has proposed that these regions are central to mentalising and animacy (e.g. ^13,16,18^). As our localisers did not reliably elicit mPFC or rmFG response in this participant cohort, we created ROI from coordinates in the original localiser paper.^50^ It is possible that our “generic” ROIs failed to capture individuals’ peak mentalizing voxels across these regions.

However, mPFC and rmFG activation clearly emerges in many of our group whole brain contrasts. Precuneus clusters in our localizer and whole brain contrasts were large and the peak cluster from the localiser was more inferior and lateral than the peak clusters in the whole brain contrasts. Last, it is also possible that our localisers produced coordinates for offline social cognition or mentalizing and not for online social cognition.^51^ Thus, it seems likely that mPFC, rmFG, and Precuneus do likely play a role in mentalizing in our study, but were not well captured by our chosen ROI coordinates.

We also explored the response profile of a region in the pSTS that is sensitive to interactive information in observed dyadic social interactions.^52^ This region is nearby, but distinct from the TPJ, and might plausibly discriminate between game partners. Response in the pSTS discriminated between game partners both during game play and during the video preceding each game series. This was somewhat surprising as the pSTS is largely responsive to the perceptual features of interactions, particularly biological motion.^53,54^ In our design, there were no social perceptual features to process during game play as players observed the same visual stimuli during game play across all four conditions. This suggests that perhaps top-down knowledge cues may be more influential in this region than previously thought. We further explored this data by testing our linear human-likeness and socialness models on the pSTS data from video 1 and gameplay. Both models were significant, but in this case the social model was numerically stronger during both gameplay (rpSTS only) and video 1 (bilateral pSTS). The pSTS has been implicated as a part of the social cognition and mentalising networks and has previously been shown to integrate both perceptual and social features.^52,55–57^ The pSTS also responds strongly to social interactions between non-human agents such as moving shapes and dots of light that mimic social scenarios (e.g. ^55,57,58^), and do so even more strongly when participants are led to believe an object is animate versus inanimate.^36,59^ One possibility is that because participants were engaging in a real-time interaction in our study, the pSTS was more strongly driven by the social features of game partners rather than their visual features. When motion and visual cues to humanness conflict or are not reliably aligned with more top-down attributions of socialness, the more superior regions in the pSTS may prioritise top-down knowledge cues to humanness in social interactions.

While our neuroimaging and behavioural results indicate a linear effect of human likeness and socialness across conditions, pairwise comparisons from our ROIs also show that the human partner is perceived significantly differently from all others game partners. While this result is perhaps unsurprising, it suggests that a uniquely human factor still differentiates people from animate non-human entities, even when they are quite ‘human like’ in appearance or behaviour. This result has been reported previously,^36,42^ and is consistent with the idea that the mentalizing system may be best tuned to human actors and human social cues. It is possible with advancing technology and design that the line between robots and humans may blur, and mentalising regions will become increasingly recruited.

One surprise in our results is that game play did not drive responses in mentalizing regions above baseline. Our expectation, based on prior,^26,43^ was that this task would indeed drive engagement of the mentalizing network, at least for the human partner, above baseline. Previous studies^26,27^ found negative activation to the computer condition and to the non-android robots in mentalizing ROIs, but above-baseline response to the human partner. One possibility here is that our task was particularly demanding, requiring not only mentalizing but also analysing and remembering strategies for each opponent. Thus, we think it is likely that the negative responses seen in our results are a result of most mentalizing regions being part of, or close to, the default mode network, which tends to deactivate during difficult or demanding tasks.^60^ If true, an active baseline^61^ such as the computer condition, might be a better baseline for computing response differences across the experimental conditions. Despite the negative response in mentalizing regions, the between-condition differences, evident in mentalizing regions across both ROI and whole-brain analyses, are still interpretable.

As with previous studies, and unbeknownst to the participants, we controlled wins and losses amongst game partners so our findings could not be explained by winning or losing more to any one partner. Participants’ ratings of success and strategy against each of the 4 game partners did not significantly differ, suggesting that they accurately perceived their own performance, including that their strategy did not work any more efficiently for one partner than another, similar to previous findings.^26^ Therefore, it is unlikely that our findings are due to perceived differences in difficulty in playing each partner. Employing a strategic approach to the game likely relates to thinking about the mind of the other player, and thus to activity in the mentalizing network. As a result, participants in this study may have reduced their mentalizing about game partners as they found that their strategies were not working. Future studies might look at manipulating wins and losses or alter initial briefing instructions to create different impressions of each game partner’s fun and competitiveness in order to explore the extent to which socialness can be manipulated to influence mind attribution toward robots.

### Theoretical implications

Our results support growing evidence emerging from the intersection of social robotics and social neuroscience that multiple routes exist to non-human agents being perceived as “like-me”,^36,42^ including not only a human-like appearance or motion profile, but also being perceived as ‘social’ based on behaviours or background knowledge about a robot. Significant R&D investment continues to fuel the development of socially interactive robots with whom human users can intuitively and effectively collaborate, which often attempt to capture as much human-likeness as possible while also avoiding the uncanny valley.^62–67^ However, the extent to which an agent is perceived as “like-me” extends beyond physical form, capabilities, and movement, and growing evidence supports that prior knowledge about and the perceived socialness of a robot may more strongly influence their reception (and people’s ability to collaborate or cooperate with them in an intuitive manner) in social settings.^41,44,68–73^

A few neuroimaging studies have investigated how these top-down knowledge cues and bottom-up stimulus cues influence perceptions of animacy and the flexibility of our social cognitive system. One study found that stimulus cues overrode knowledge cues to animacy ^74^; whereas, others found the inverse, knowledge, not stimulus, cues more strongly influenced animacy perception.^42,75^ Yet, a key mentalizing region (rTPJ) was most sensitive when *both* stimulus and knowledge cues to animacy were presented compared to when only one (or none) of those cues were present.^36^ These various findings are likely influenced by the type of task and cues used, and our study adds to the narrative that top-down knowledge-based cues of socialness can be just as, if not more, powerful for driving mind attribution during social interactions with artificial agents than bottom-up visual cues to human-likeness alone.

Therefore, physical features denoting human-likeness may not be the most important consideration for those designing socially engaging robots, and instead a reorientation toward an emphasis on socialness may be more fruitful for fostering social behaviours and attitudes toward robots. Ultimately, our findings set the stage for future work to disentangle not only which physical and social features play the most important roles in mind attribution to artificial agents, but also how ongoing experience with such agents changes and develops such perceptions.

### Concluding thoughts

Our primary findings confirm previous research that human-likeness plays an important role in the attribution of mind to robots. However, our exploratory analyses suggest that the perceived socialness of a robot also plays an equally, if not more important role than physical features denoting human-likeness in mind attribution. Incorporating knowledge- or experience-based social cues and features into robots who are designed to engage human users on a social level has the potential to increase user engagement and interest for more lasting and higher quality relationships with our robotic partners.

## STAR METHODS

| REAGENT or RESOURCE | SOURCE | IDENTIFIER |
| --- | --- | --- |
| **Deposited Data** |  |  |
| Data supporting the main findings<sup>1</sup> | OpenNeuro |  |
| Group level whole brain results | Neurovault | https://neurovault.org/collections/EERVYAPX/ |
| Behavioural data & stimuli examples | OSF |  |
| Preregistration | AsPredicted.org | https://aspredicted.org/CBG_ZPG |
| **Software & Algorithms** |  |  |
| fMRI analyses | Matlab 2018a, MathWorks Inc<br>https://it.mathworks.com/products/matlab.html | RRID:SCR_001622 |
| fMRI analyses | Statistical Parametric Mapping 12 (SPM12)<br>https://www.fil.ion.ucl.ac.uk/spm/ | RRID:SCR_007037 |
| Visual Presentation of Main Task and to create robot actions | Python software foundation<br>http://www.python.org/ | RRID:SCR_008394 |
| Visual Presentation of localiser (TOM) | Psychopy<br>https://www.psychopy.org/ | RRID:SCR_006571 |
| Visual presentation of localiser (INT) | PsychToolBox 3<br>http://psychtoolbox.org/ | RRID:SCR_002881 |
| R Project for Statistical Computing (for figs) | The R Foundation<br>http://www.r-project.org/ | RRID:SCR_001905 |

<sup>1</sup> Bangor neuroimaging data are available for analysis through request due to BIDS conversion difficulties with Phillips multi-echo data

|  |  |  |
| --- | --- | --- |
| Code for robot introduction & main experiment | GitHub | https://github.com/chaudhuryB |
| Code for analyses | upon request |  |

### Human subjects

Due to the availability of scanning resources, participants were recruited from 2 sites: (i) the greater Glasgow area (Scotland, UK); and (ii) the greater Bangor area (Wales, UK). Glasgow participants completed the study at the Centre for Cognitive Neuroimaging (CCNi) at the University of Glasgow, while Bangor participants completed the study at the Bangor Imaging Unit (BIU) at Bangor University.

Twenty right-handed males (mean age = 20.95 years; SD = 1.82; range = 19-26) participated from the greater Bangor area and 24 right-handed males (mean age = 22.45 years; SD = 3.63; range = 18-32) participated from Glasgow. Only males were recruited, consistent with previous studies in this area,^26,27^ in order to control for any potential effects of gender on mentalizing.^78^ Two participants withdrew from the study due to claustrophobia (1 subject from each site). The final fMRI participant sample included a total of 42 participants (mean age = 21.74 years; SD = 3.03; range = 18-32).

All participants reported normal or corrected-to-normal vision, no history of neurological or psychiatric disorders, and were right-handed as confirmed on the Edinburgh Handedness Questionnaire^79^; mean = 1.48, sd = .34).

All participants reported low familiarity with robots. Median engagement with robots in daily life (measured from 1 (never) to 7 (daily)) was 2 (IQR 1). Median number of robot-themed movies or TV shows seen was 4 (IQR 1) out of the 14 listed (Riek et al, 2011).^80^

All participants provided written informed consent prior to their involvement and received monetary compensation for study participation (£12/hour). All study procedures were approved by the respective university ethics boards: (i) Bangor University (Approval no. 2019-16639) and (ii) Glasgow University (Approval no. 300180110).

Study site was a significantly different between subjects factor in rmANOVA for rTPJ, rmFG & Precuneus but when we ran site separately for each of those ROIs, the results did not differ from the combined group or change the outcome, therefore, both sites were kept together in the main tables above. Please see our OSF for details on the results from the separate groups. Further, we ran site as a covariate of no-interest in our model estimation and did not find differences in our whole brain data; therefore, the sites were subsequently analysed and reported together (please see our OSF for more details).

### Experimental Design

We designed a Rock-Paper-Scissors (RPS) task similar to a previous study,^26^ and followed a similar briefing procedure.^26,27^ RPS was chosen for its familiarity across ages and cultures, and ease of rule learning. Previous studies have shown this game to engage mentalizing regions when played against human and non-human partners.^26,29,43^

Participants saw videos of their respective game partners before and after each 3-game series (see Fig 2). Each video was unique and all participants saw the same set of videos. The button press for rock, paper, and scissors were assigned randomly across participants.

In-line with previous designs,^26,27,43^ participants were told that they were playing a live game and viewing their game-partners through a live video feed, but in reality, neither the remote practice nor the in-scanner games (described below) were live. All videos were pre-recorded and designed to give the impression of a live game. Wins and losses were controlled across the four conditions so that each participant won 10 rounds and lost 10 rounds against each partner. The order in which participants played partners was pseudo-randomized across four 8-minute functional runs.

To give the impression of a live game, participants met all game partners in person in the “game room” and played one truly live, in-person, round of rock-paper-scissors with each partner. They went to the imaging suite to play a “live” practice round of RPS with their partners via the “video feed”. This practice round served to familiarise participants with the game and practice pushing the buttons to register their answer with the correct timing. Participants played each partner twice in each practice round and could complete up to 3 practice rounds (24 total games) to ensure they understood the game before entering the scanner. All participants demonstrated understanding of the game and button presses by the 3^rd^ practice round.

Participants then completed the fMRI task, playing the same RPS game. Each fMRI run contained 5 rounds with each of the 4 partners (20 rounds per partner across all 4 runs), pseudorandomized across participants. In total, each participant completed four, 8-minute RPS runs. After the scan, participants completed several questionnaires (listed below) on a laptop and were then debriefed. The debriefing unveiled the study deception (that the various game partners were pre-recorded, not live, and that all partners used the same random algorithm and were not independently controlled). Both the practice round and game in the scanner were programmed in Python 3.7 and run from the command line (see STAR Methods Table).

## MATERIALS

### MRI Parameters, pre-processing, & GLM estimation

#### fMRI data acquisition & Pre-processing

At both data collection sites (CCNI and BIU), stimuli were projected onto a mirror from a projector located behind the scanner. Responses were recorded with an MRI-compatible keypad.

A dual-echo EPI sequence was used to improve signal-to-noise ratio (SNR) in frontal and temporal regions.^81^ All structural and functional sequence parameters are detailed in Supplemental Tables 1 & 2.

Data pre-processing was carried out in SPM12 (Wellcome Trust Centre for Neuroimaging, London) implemented in Matlab 2018a (Mathworks, Natick, MA, USA). Pre-processing consisted of standard SPM12 defaults for slice time correction, realignment and re-slicing, co-registration, unified segmentation & normalisation, and smoothing; except for a 6mm FWHM Gaussian smoothing kernel. All analyses were performed in normalized MNI space. Block durations and onsets for each of the 4 experimental conditions during Video 1, the RPS game, and Video 2 were modelled by convolving the hemodynamic response function and with a high pass filter of 128s. Head motion parameters were modelled as nuisance regressors. Functional scans provided whole brain coverage.

### ROI Creation & Analyses

Our choice of ROIs was informed by previous studies^26,27^; however, ROI placement was based on peak activation from the independent localizers (see Fig S2 & Table S3). Participants undertook two passive-viewing tasks to help identify brain regions of interest after playing the RPS game. Localizer 1 was a short-animated film (‘Partly Cloudy’; Pixar Animation Studios, 2009) used to localize bilateral TPJ, bilateral mFG, and Precuneus with the mentalizing > pain contrast. Neither Medial Prefrontal Cortex (mPFC) nor rmFG activation appeared as expected in Localiser 1, therefore, we used mPFC & rmFG coordinates from the original localiser paper^50^ and created 6mm spheres around those coordinates. Localizer 2 was a point-light figure social interaction task^52,57^ to localize bilateral pSTS with the interaction > scrambled contrast (i.e., two human point light figures interacting vs. scrambled dot motion).

We used a control ROI (V1/BA17) from the WFU PickAtlas^82^ as a form of verification that activity differences seen between conditions during game play was not attributable to non-specific whole brain activation differences. In other words, we would not expect differences between conditions in V1 activity during game play, as participants saw the same set-up across all conditions, and this control ROI allowed us to evaluate this possibility.

Briefly, group-constrained, subject specific ROIs were created like the methods described elsewhere^52^ using an uncorrected height threshold of p < .0001. This protocol creates subject-specific ROIs using a leave-one-subject-out iterative process so that ROIs for individuals are based on independent data. ROIs were created based on group activation from the localizer tasks defined by intersecting a 6mm sphere with the cluster peak (i.e. highest voxel t-value) in each of the regions named above. That subject specific search space was then applied to the participant’s individual data to create the final ROIs. Percent signal change was extracted from ROIs using in-house scripts in Matlab 2018a and the MarsBar toolbox.

None of the ROIs overlapped. Both right and left TPJ were slightly shifted so the entire sphere was within the boundaries of the brain; all other ROIs created from the localisers remained true to the peak activation. Please refer to Supplementary Figures for all ROI coordinates.

#### Pre-Registered

Repeated measures ANOVAs were run for each ROI to assess the effect of game-partner and pairwise comparisons were run only if a main effect of game-partner was found. We assessed the linear effect of human-likeness using a linear repeated contrast in a within-subject ANOVA, which compares means across the different levels of the independent variable according to the following order: computer < mechanoid < humanoid < human.

#### Exploratory

Ratings results from the *Fun*, *Competitiveness*, and *Sympathy* questions in the Debrief, suggested swapping the robot orders in the linear model (see below). As an exploratory analysis, we ran a linear repeated contrast in a within-subject ANOVA to compare means across different levels of the independent variable according to the following order based on socialness ratings: computer < humanoid < mechanoid < human.

Additionally, we assessed whether the pSTS would show a linear pattern based on human-likeness or socialness during game play and whilst watching the video introduction which preceded each round.

### Whole brain analyses

#### Pre-Registered

A GLM comprising the four conditions (CP = Computer Partner, MR = Mechanoid Robot, HR= Humanoid Robot, HP = Human Partner) was specified for each participant. Simple contrasts were compared against: (1) HP > CP, (2) HR > CP, (3) MR > CP, (4) HR > MR, (5) HP > HR. Based on previous findings (Krach et al, 2010) and our hypothesis, we expected to see a linear increase in neural activity based on human-likeness of agent. To evaluate this, we calculated a parametric modulation of gameplay partner (actual model weights used: CP = -3, MR = -1, HR = 1, HP = 3). For the second level group analyses, we used a FWE-corrected threshold (p_uncorr_ < 0.001) and a minimum cluster size (k = 100).

#### Exploratory

While not pre-registered, we also included the following simple contrasts: (6) HP > MR, (7) MR > HR. We also calculated the parametric modulation of gameplay partners based on socialness (actual model weights used: CP = -3, HR = -1, MR = 1, HP = 3).

### Behavioural Measures

#### Debrief Questions

##### Pre-Registered

After scanning, participants answered questions about their experience of the study using FormR.^83^ Participants rated responses to the following questions on a scale from 0-10: (i) how well they were able to adopt an efficient *strategy* against each partner, (ii) how *successful* they were against each partner, (iii) how much *fun* it was to play each partner, (iv) how much *sympathy* they had for each partner when they lost, and then each partner’s (v) *competitiveness,* and (v) *intelligence*. As pre-registered, rmANOVAs were run on each question to assess the effect of agent. Pairwise comparisons between agents were run only if an agent effect was identified. We assessed the linear effect of human-likeness using a linear repeated contrast in a within-subject ANOVA, which compares means across different levels of the independent variable.

##### Exploratory

Furthermore, based on participant-reported perceptions of socialness of the individual agents, we ran an exploratory (not pre-registered) linear repeated contrast in a within-subjects ANOVA that reversed the order of the robots in the linear model.

#### Inclusion of Others and Self (IOS)

The Inclusion of Others and Self (IOS) is a measure of closeness and interconnectedness between two individuals.^84^ A series of 7 increasingly overlapping circles are presented to the participant on paper. Each pair of circles contains the word “self” in one circle and “other” in the other circle. Participants are then asked to choose which circle represents their relationship to the agent in question. We asked participants to show which set of overlapping circles best describes the following agents: (1) computer, (2) mechanoid robot, (3) humanoid robot, (4) a human stranger, (5) the human from the experiment (LEJ), and (6) a close friend. Non-robot items were included for comparison to determine where the robot stood relative to other people in the participant’s lives. The IOS provides another way to address the participant’s view of their relationship to various humans and robots. Responses from the paper and pencil format of the IOS were recorded onto a 7-point scale from 1 (no overlap) to 7 (nearly complete overlap). As pre-registered, rmANOVA was run to assess the effect of agent and pairwise comparisons were run only if an effect of agent was found.

## Supplementary Data

See Supplementary Data section for more detail.

## Acknowledgements

This work has received funding from the European Research Council under the European Union’s Horizon 2020 research and innovation programme (Grant Agreement numbers: 677270 (Social Robots) & 716974: Becoming Social)).

The authors thank Nikolas Vitsakis, Kiara Jackson, and Jacynth Grundy for assistance with data collection and Julia Landsiedel for programming advice.

## Author Contributions

Conceptualisation: LEJ, ESC, KK; Methodology: LEJ, ESC, KK, BC; Formal Analysis: LEJ; Investigation: LEJ, BC, SAA; Writing - Original Draft : LEJ, ESC, KK; Writing - Review & Editing: LEJ, ESC, KK, BC, SAA; Supervision: ESC & KK; Funding Acquisition: ESC & KK.

