## Supplemental Information for "Beyond human-likeness: Socialness is more influential when attributing mental states to robots"

Supplemental Materials

| Table S1. T1 acquisition parameters by site | | |
|---|---|---|
| | Site 1 | Site 2 |
| Repetition time (ms) | 23 | 18 |
| Echo time (ms) | 3.0 | 3.5 |
| Acquired voxel size (mm) | 1 x 1 x 2 | 1 x 1 x 2 |
| Reconstructed voxel size (mm) | 1 x 1 x 1 | 1 x 1 x 1 |
| Number of slices (contiguous axial) | 176 | 192 |
| Flip angle | 9° | 8° |
| Field of view (mm) | 256 x 256 | 256 x 256 |
| Acquisition matrix size | 256 x 220 | 256 x 220 |
| Reconstruction matrix | 256 | 256 |
| Total acquisition time | 6 min | 6 min |
| Pulse sequence | T1 | T1 |
| 3T MRI system | Siemens TrioTim | Philips Ingenia Elition |
| Site | University of Glasgow | Bangor University |
| Both sites used a gradient echo, multi-shot turbo field echo pulse sequence. | | |

| Table S2. BOLD acquisition parameters by site | | |
|---|---|---|
| | Site 1 | Site 2 |
| Repetition time (ms) | 2000 | 2000 |
| Echo time (ms) | 13 / 31 | 12 / 31 |
| Voxel size (mm) | 2.75 x 2.75 x 4 | 2.75 x 2.75 x 4 |
| Number of slices | 32 | 32 |
| Flip angle | 85° | 85° |
| Field of view (mm) | 220 x 220 | 220 x 220 |
| Matrix size | 80 x 80 | 80 x 80 |
| Slice acquisition | Ascending | Ascending |
| Phase encoding | Anterior-posterior | Anterior-posterior |
| Acquisition orientation | 45° from AC-PC | 45° from AC-PC |
| Number of volumes (RPS)* | 239 | 240 |
| Number of volumes (ToM) | 174 | 174 |
| Number of volumes (INT)* | 76 | 72 |
| Pulse sequence | dual-echo EPI | dual-echo EPI |
| Parallel imaging parameters | GRAPPA accel. factor (3) | SENSE factor (3) |
| Head coil | 32-channel | 32-channel |
| 3T MRI system | Siemens TrioTim | Philips Ingenia Elition |
| Site | University of Glasgow | Bangor University |

RPS = Rock Paper Scissors task (main task); ToM = Theory of Mind localizer task (Partly Cloudy movie; Jacoby et al., 2018); INT = Interaction Localizer task (Walbrin & Koldewyn, 2018); Number of volumes for each functional scan were trimmed for analyses so that there were the same number of volumes per site resulting in: (1) 239 volumes for the RPS task, (2) 72 volumes for the Interaction Task (INT).

Table S3. Whole-brain activation peaks with their localization

| | | Coordinates | | | Peak | | Cluster-level | |
|---|---|---|---|---|---|---|---|---|
| | | x | y | z | t-value | k | $p_{uncorr}$ | $p_{FWE}$ |
| **ToM > Pain** | | | | | | | | |
| - | Precuneus | 16 | -56 | 22 | 8.30 | 2524 | <0.001 | <0.001 |
| R | Temporal-Parietal Junction* | 48 | -65 | 32 | 8.50 | 1217 | <0.001 | <0.001 |
| L | Temporal-Parietal Junction | -46 | -62 | 38 | 6.90 | 923 | <0.001 | <0.001 |
| L | Middle Frontal Gyrus | -36 | 8 | 54 | 5.14 | 145 | 0.003 | 0.043 |
| **Interaction > Non-Interaction** | | | | | | | | |
| R | Posterior Superior Temp Sulcus | 54 | -48 | 18 | 4.92 | 448 | <0.001 | <0.001 |
| L | Posterior Superior Temp Sulcus* | -56 | -54 | 22 | 5.29 | 158 | 0.004 | 0.048 |
| **Additional ROIs from other sources** | | | | | Coordinate Source | | | |
| | Dorsal Medial Prefrontal Cortex | -10 | 56 | 32 | Jacoby et al. (2016) | | | |
| | Ventral Medial Prefrontal Cortex | 2 | 54 | -12 | Jacoby et al. (2016) | | | |
| R | Middle Frontal Gyrus | 24 | 28 | 40 | Jacoby et al. (2016) | | | |
| R | V1 (BA 17) | | | | WFU PickAtlas | | | |
| L | V1 (BA 17) | | | | WFU PickAtlas | | | |

Significance level and size of activation cluster for the primary contrast (both $p_{uncorr}$ and $p_{FWEcorr}$ reported; minimal cluster size, k ≥ 0). Coordinates reported in MNI space. L = Left; R = Right. ToM Localizer uses a short film by Pixar ("Partly Cloudy") to localize mentalizing regions of the brain (Jacoby et al. 2016). As our ToM localizer did not provide mPFC or rmFG coordinates, we created 6mm spheres in Marsbar for those ROIs based on coordinates from Jacoby et al. (2016). rTPJ (true peak coordinates: 50 -68 32) & lpSTS (true peak coordinates: -62 -54 22) coordinates reported in the table above were shifted from true peak so that spheres remained within the brain boundaries.

Table S4. Whole Brain activation peaks with their localization for RPS task

| | | MNI Coordinates (mm) | | | Peak t-value | Cluster Extent (voxels) | Cluster-level | |
|---|---|---|---|---|---|---|---|---|
| | | x | y | z | | k | $p_{uncorr}$ | $p_{FWE}$ |
| **Mechanoid > Computer** | | | | | | | | |
| | - | - | - | - | - | - | ns | ns |
| **Humanoid > Computer** | | | | | | | | |
| | - | - | - | - | - | - | ns | ns |
| **Human > Computer** | | | | | | | | |
| - | Medial Prefrontal Cortex | 4 | 48 | 24 | 5.95 | 1264 | <0.001 | <0.001 |
| R | Temporal-Parietal Junction | 50 | -46 | 20 | 5.42 | 864 | <0.001 | <0.001 |
| - | Thalamus | 6 | 8 | 2 | 5.23 | 619 | <0.001 | 0.001 |
| - | Superior Medial Prefrontal Cortex | -2 | 28 | 58 | 5.11 | 632 | <0.001 | <0.001 |
| R | Posterior Superior Temporal Sulcus | 54 | -20 | -14 | 5.07 | 330 | 0.001 | 0.005 |
| | Precuneus | -2 | -50 | 30 | 4.82 | 859 | <0.001 | <0.001 |
| L | Middle Frontal Gyrus | -42 | 14 | 50 | 4.65 | 318 | 0.001 | 0.006 |
| R | Middle Frontal Gyrus | 40 | 12 | 52 | 4.58 | 285 | 0.001 | 0.010 |
| L | Temporal-Parietal Junction | -54 | -50 | 26 | 4.56 | 390 | <0.001 | 0.002 |
| L | Cerebellum | -14 | -82 | -32 | 4.33 | 252 | 0.002 | 0.018 |
| **Human > Humanoid** | | | | | | | | |
| R | Temporal-Parietal Junction | 46 | -54 | 28 | 7.24 | 1556 | <0.001 | <0.001 |
| R | Pars Triangularis (IFG) | 48 | 26 | 18 | 7.06 | 3731 | <0.001 | <0.001 |
| R | Posterior Superior Temporal Sulcus | 60 | -22 | -12 | 6.37 | 1238 | <0.001 | <0.001 |
| - | Precuneus | 4 | -60 | 34 | 5.65 | 1361 | <0.001 | <0.001 |
| - | Cerebellum Crus 1 | -20 | -82 | -30 | 5.18 | 600 | <0.001 | <0.001 |

| | | | | | | | | |
|---|---|---|---|---|---|---|---|---|
| R | Middle Frontal Gyrus | 34 | 32 | 44 | 4.83 | 1045 | <0.001 | <0.001 |
| - | Nucleus Accumbens | 8 | 10 | -14 | 4.78 | 222 | 0.004 | 0.031 |
| L | Temporal-Parietal Junction | -48 | -48 | 30 | 4.39 | 428 | <0.001 | 0.001 |
| Human > Mechanoid | | | | | | | | |
| R | Temporal-Parietal Junction | 48 | -44 | 26 | 5.20 | 907 | <0.001 | <0.001 |
| L | Cerebellum Crus 1 | -20 | -82 | -30 | 5.18 | 261 | 0.002 | 0.018 |
| R | Precuneus | 6 | -56 | 32 | 4.90 | 365 | <0.001 | 0.004 |
| R | Posterior Superior Temporal Sulcus | 52 | -20 | -8 | 4.21 | 212 | 0.005 | 0.039 |
| Humanoid > Mechanoid | | | | | | | | |
| - | - | - | - | - | - | - | - | - |
| Mechanoid > Humanoid | | | | | | | | |
| | Nucleus Accumbens | -4 | 10 | -10 | 4.91 | 313 | <0.001 | 0.003 |
| Human-likeness (CP > MR > HR > HP) | | | | | | | | |
| - | Medial Prefrontal Cortex | 2 | 48 | 24 | 5.46 | 483 | <0.001 | <0.001 |
| R | Temporal-Parietal Junction | 50 | -46 | 22 | 5.27 | 778 | <0.001 | <0.001 |
| - | Superior Medial Prefrontal Cortex | 0 | 32 | 54 | 4.72 | 453 | <0.001 | 0.001 |
| - | Precuneus | 12 | -50 | 42 | 4.63 | 616 | <0.001 | <0.001 |
| - | Nucleus Accumbens | 6 | 8 | 2 | 4.63 | 242 | 0.002 | 0.018 |
| R | Middle Frontal Gyrus | 34 | 54 | 4 | 4.53 | 188 | 0.005 | 0.048 |
| L | Middle Frontal Gyrus | -46 | 14 | 44 | 4.22 | 225 | 0.003 | 0.025 |
| Socialness (CP > HR > MR > HP) | | | | | | | | |
| - | Medial Prefrontal Cortex | -6 | 44 | 18 | 6.12 | 1727 | <0.001 | <0.001 |
| R | Temporal-Parietal Junction | 50 | -50 | 18 | 5.52 | 859 | <0.001 | <0.001 |
| - | Nucleus Accumbens | 6 | 8 | 2 | 5.40 | 745 | <0.001 | <0.001 |
| R | Posterior Superior Temporal Sulcus | 56 | -28 | -12 | 5.30 | 403 | <0.001 | 0.002 |
| L | Posterior Superior Temporal Sulcus | -56 | -22 | -10 | 5.26 | 242 | 0.002 | 0.022 |

| | | | | | | | | |
|---|---|---|---|---|---|---|---|---|
| L | Middle Frontal Gyrus | -24 | 54 | 18 | 5.16 | 270 | 0.002 | 0.014 |
| L | Supplementary Motor Area | -2 | 26 | 58 | 5.14 | 819 | <0.001 | <0.001 |
| - | Precuneus | 12 | -52 | 42 | 4.86 | 954 | <0.001 | <0.001 |
| L | Middle Frontal Gyrus | -42 | 14 | 50 | 4.82 | 332 | 0.001 | 0.006 |
| R | Pars Triangularis | 46 | 24 | 14 | 4.74 | 618 | <0.001 | <0.001 |
| L | Temporal-Parietal Junction | -58 | -50 | 28 | 4.64 | 506 | <0.001 | <0.001 |
| R | Middle Frontal Gyrus | 40 | 12 | 52 | 4.57 | 407 | <0.001 | 0.002 |
| L | Cerebellum Crus 2 | -12 | -82 | -44 | 4.45 | 285 | 0.001 | 0.011 |

Significance level and size of activation cluster for each of the contrasts and the parametric modulation listed above (both $p_{uncorr\ and}$ $p_{FWEcorr}$ reported; no minimal cluster threshold). Coordinates reported in MNI space.

Table s5. rmANOVAs for PSC in each ROI

| | | Within-Subjects Effects | | | | Pairwise Comparisons: P-value / Cohen's d | | | | Within-Subjects Contrasts (Linear) H: Humanness model S: Socialness model | | | | |
|---|---|---|---|---|---|---|---|---|---|---|---|---|---|---|
| | Mean ± SD | *df* | *F* | p | $\eta_p^2$ | CP | MR | HR | HP | | *df* | *F* | p | $\eta_p^2$ |
| **rTPJ**[§] | | | | | | | | | | | | | | |
| Computer | -0.15 ± 0.21 | 2.5, 103.83 | 6.78 | <0.001 | 0.14 | | 0.09 | 0.14 | -0.14 | H | 1, 41 | 3.67 | 0.063 | 0.08 |
| Mechanoid | -0.17 ± 0.23 | | | | | ns | | 0.04 | -0.22 | | | | | |
| Humanoid | -0.18 ± 0.23 | | | | | ns | ns | | -0.26 | S | 1, 41 | 4.53 | 0.039 | 0.10 |
| Human | -0.12 ± 0.23 | | | | | ns | ** | *** | | | | | | |
| **lTPJ** | | | | | | | | | | | | | | |
| Computer | -0.03 ± 0.23 | 3, 123 | 7.60 | <0.001 | 0.16 | | -0.09 | 0 | -0.37 | H | 1, 41 | 7.60 | <0.001 | 0.16 |
| Mechanoid | -0.01 ± 0.21 | | | | | ns | | 0.10 | -0.29 | | | | | |
| Humanoid | -0.03 ± 0.21 | | | | | ns | ns | | -0.39 | S | 1, 41 | 12.68 | <0.001 | 0.24 |
| Human | 0.05 ± 0.20 | | | | | *** | ** | ** | | | | | | |
| **rmFG**[§] | | | | | | | | | | | | | | |
| Computer | -0.12 ± 0.21 | 2.56, 104.84 | 2.28 | 0.092 | 0.05 | | | | | | | | | |
| Mechanoid | -0.14 ± 0.20 | | | | | | | | | | | | | |
| Humanoid | -0.15 ± 0.21 | | | | | | | | | | | | | |
| Human | -0.11 ± 0.19 | | | | | | | | | | | | | |
| **lmFG**[§] | | | | | | | | | | | | | | |
| Computer | 0.00 ± 0.17 | 2.27, 92.92 | 3.17 | 0.040 | 0.07 | | -0.17 | 0 | -0.28 | H | 1, 41 | 4.62 | 0.038 | 0.10 |
| Mechanoid | 0.03 ± 0.18 | | | | | ns | | 0.17 | -0.11 | | | | | |
| Humanoid | 0.00 ± 0.18 | | | | | ns | ns | | -0.27 | S | 1, 41 | 7.11 | 0.011 | 0.15 |
| Human | 0.05 ± 0.19 | | | | | ns | ns | ns | | | | | | |
| **Precuneus** | | | | | | | | | | | | | | |

| | | | | | | | | | | | | | | |
|---|---|---|---|---|---|---|---|---|---|---|---|---|---|---|
| Computer | -0.18 ± 0.17 | 3, 123 | 1.04 | 0.376 | 0.03 | | | | | | | | | |
| Mechanoid | -0.17 ± 0.19 | | | | | | | | | | | | | |
| Humanoid | -0.19 ± 0.18 | | | | | | | | | | | | | |
| Human | -0.16 ± 0.20 | | | | | | | | | | | | | |
| **dmPFC** | | | | | | | | | | | | | | |
| Computer | -0.22 ± 0.21 | 3, 123 | 2.19 | 0.092 | 0.05 | | | | | | | | | |
| Mechanoid | -0.20 ± 0.20 | | | | | | | | | | | | | |
| Humanoid | -0.24 ± 0.22 | | | | | | | | | | | | | |
| Human | -0.20 ± 0.20 | | | | | | | | | | | | | |
| **vmPFC** | | | | | | | | | | | | | | |
| Computer | -0.50 ± 0.44 | 3, 123 | 2.16 | 0.097 | 0.05 | | | | | | | | | |
| Mechanoid | -0.49 ± 0.50 | | | | | | | | | | | | | |
| Humanoid | -0.55 ± 0.51 | | | | | | | | | | | | | |
| Human | -0.47 ± 0.43 | | | | | | | | | | | | | |
| **rpSTS** | | | | | | | | | | | | | | |
| Computer | -0.02 ± 0.25 | 3, 123 | 12.39 | <0.001 | 0.23 | | -0.08 | 0 | -0.38 | H | 1, 41 | 18.87 | <0.001 | 0.315 |
| Mechanoid | 0.00 ± 0.25 | | | | | ns | | 0.08 | -0.30 | | | | | |
| Humanoid | -0.02 ± 0.25 | | | | | ns | ns | | -0.38 | S | 1,41 | 25.27 | <0.001 | 0.381 |
| Human | 0.07 ± 0.22 | | | | | *** | ** | *** | | | | | | |
| **rpSTS** (Video 1) | | | | | | | | | | | | | | |
| Computer | 0.07 ± 0.14 | 3, 123 | 29.40 | <0.001 | 0.42 | | -0.73 | -0.54 | -1.40 | H | 1, 38 | 52.30 | <0.001 | 0.561 |
| Mechanoid | 0.20 ± 0.21 | | | | | *** | | 0.20 | -0.57 | | | | | |
| Humanoid | 0.16 ± 0.19 | | | | | ns | 0.029 | | -0.80 | S | 1, 41 | 77.98 | <0.001 | 0.655 |
| Human | 0.32 ± 0.21 | | | | | *** | *** | *** | | | | | | |
| **lpSTS** | | | | | | | | | | | | | | |
| Computer | -0.11 ± 0.26 | 3, 123 | 6.96 | <0.001 | 0.15 | | -0.08 | 0 | 0.91 | H | 1,41 | 16.33 | <0.001 | 0.285 |
| Mechanoid | -0.09 ± 0.26 | | | | | ns | | 0.07 | 0.98 | | | | | |
| Humanoid | -0.11 ± 0.29 | | | | | ns | ns | | 0.86 | S | 1, 41 | 16.22 | <0.001 | 0.283 |

| | | | | | | | | | | | | | | |
|---|---|---|---|---|---|---|---|---|---|---|---|---|---|---|
| Human | -0.35 ± 0.27 | | | | | *** | * | * | | | | | | |
| **lpSTS** (Video1) | | | | | | | | | | | | | | |
| Computer | 0.04 ± 0.20 | 3, 123 | 13.26 | <0.001 | 0.24 | | -0.32 | -0.20 | -0.79 | H | 1, 41 | 22.64 | <0.001 | 0.356 |
| Mechanoid | 0.10 ± 0.18 | | | | | 0.056 | | 0.11 | -0.50 | | | | | |
| Humanoid | 0.08 ± 0.20 | | | | | ns | ns | | -0.58 | S | 1, 41 | 31.27 | <0.001 | 0.433 |
| Human | 0.19 ± 0.18 | | | | | *** | ** | *** | | | | | | |
| **rV1[§] (BA17)** | | | | | | | | | | | | | | |
| Computer | 0.20 ± 0.39 | 2.42, 91.93 | 1.10 | 0.341 | 0.03 | | | | | | | | | |
| Mechanoid | 0.20 ± 0.34 | | | | | | | | | | | | | |
| Humanoid | 0.19 ± 0.37 | | | | | | | | | | | | | |
| Human | 0.25 ± 0.36 | | | | | | | | | | | | | |
| **lV1[§] (BA17)** | | | | | | | | | | | | | | |
| Computer | 0.09 ± 0.27 | 2.50, 102.19 | 0.97 | 0.400 | 0.23 | | | | | | | | | |
| Mechanoid | 0.09 ± 0.27 | | | | | | | | | | | | | |
| Humanoid | 0.08 ± 0.26 | | | | | | | | | | | | | |
| Human | 0.12 ± 0.23 | | | | | | | | | | | | | |
| Note: Total N = 42. All p-values reported are Bonferroni corrected, *p<0.05, **p<0.01, ***p<0.001. [§]Greenhouse-Geisser correction. | | | | | | | | | | | | | | |

Table s6. rmANOVAs for Debrief Questions

| | | **Within-Subjects Effects** | | | | **Pairwise Comparisons** (P-value / Cohen's d) | | | | **Within-Subjects Contrasts (Linear)** H: Humanness S: Socialness | | | | |
|---|---|---|---|---|---|---|---|---|---|---|---|---|---|---|
| | **Mean ± SD** | ***df*** | ***F*** | **p** | **$\eta_p^2$** | **CP** | **MR** | **HR** | **HP** | | ***df*** | ***F*** | **p** | **$\eta_p^2$** |
| **Successful** | | | | | | | | | | | | | | |
| Computer | 6.00 ± 2.04 | 3, 123 | 0.50 | 0.685 | 0.012 | - | | | | - | | | | |
| Mechanoid | 5.83 ± 1.83 | | | | | | | | | | | | | |
| Humanoid | 5.52 ± 1.49 | | | | | | | | | | | | | |
| Human | 5.76 ± 1.78 | | | | | | | | | | | | | |
| **Strategy** | | | | | | | | | | | | | | |
| Computer | 5.24 ± 2.45 | 3, 123 | 0.32 | 0.811 | 0.008 | - | | | | - | | | | |
| Mechanoid | 5.29 ± 1.83 | | | | | | | | | | | | | |
| Humanoid | 5.36 ± 1.87 | | | | | | | | | | | | | |
| Human | 6.62 ± 1.78 | | | | | | | | | | | | | |
| **Intelligence**[§] | | | | | | | | | | | | | | |
| Computer | 5.21 ± 2.82 | 2.51, 102.91 | 12.16 | < 0.001 | 0.229 | | -0.09 | -0.34 | -0.92 | H | 1, 41 | 25.64 | <.001 | .385 |
| Mechanoid | 5.43 ± 1.96 | | | | | ns | | -0.30 | -1.02 | | | | | |
| Humanoid | 6.05 ± 2.13 | | | | | ns | ns | | -0.69 | S | 1, 41 | 14.80 | <.001 | .265 |
| Human | 7.48 ± 2.03 | | | | | *** | *** | .007 | | | | | | |
| **Fun** | | | | | | | | | | | | | | |
| Computer | 3.74 ± 2.43 | 3, 123 | 33.90 | < 0.001 | 0.453 | | -1.39 | -0.74 | -1.75 | H | 1, 41 | 60.75 | <.001 | .597 |
| Mechanoid | 7.02 ± 2.30 | | | | | *** | | 0.64 | -0.18 | | | | | |
| Humanoid | 5.52 ± 2.39 | | | | | *** | .006 | | -0.91 | S | 1, 41 | 85.74 | <.001 | .677 |
| Human | 7.38 ± 1.65 | | | | | *** | ns | *** | | | | | | |

| | | | | | | | | | | | | | | |
|---|---|---|---|---|---|---|---|---|---|---|---|---|---|---|
| **Competitive** | | | | | | | | | | | | | | |
| Computer | 3.24 ± 3.03 | 3, 123 | 17.24 | < 0.001 | 0.296 | | -1.16 | -0.68 | -115 | H | 1, 41 | 25.99 | <.001 | .388 |
| Mechanoid | 6.57 ± 2.69 | | | | | *** | | 0.47 | 0.15 | | | | | |
| Humanoid | 5.26 ± 2.87 | | | | | *** | .049 | | -0.38 | S | 1, 41 | 34.36 | <.001 | .456 |
| Human | 6.21 ± 2.68 | | | | | *** | ns | ns | | | | | | |
| **Sympathy**[§] | | | | | | | | | | | | | | |
| Computer | 0.71 ± 1.35 | 2.50, 102.58 | 58.59 | < 0.001 | 0.588 | | -1.08 | -0.98 | -2.87 | H | 1, 41 | 136.94 | <.001 | .770 |
| Mechanoid | 3.45 ± 3.32 | | | | | *** | | 0.28 | -1.11 | | | | | |
| Humanoid | 2.64 ± 2.44 | | | | | *** | ns | | -1.63 | S | 1, 41 | 149.58 | <.001 | .785 |
| Human | 6.81 ± 2.68 | | | | | *** | *** | *** | | | | | | |

Note: Total N = 42. All p-values reported are Bonferroni corrected, *p<0.05, **p<0.01, ***p<0.001. [§]Greenhouse-Geisser correction. Cohen's D for paired-sample t-test: $(x_1 - x_2) / S_{pooled}$ ($S_{pooled} = \sqrt{(S_1^2 - S_2^2) / 2}$ ).

Table S7. Inclusion of Others and Self (IOS)

| | Mean ± SD | *df* | *F* | p | $\eta p^2$ | Pairwise Comparisons | | | | | |
|---|---|---|---|---|---|---|---|---|---|---|---|
| | | | | | | P-value / Cohen's d | | | | | |
| | | | | | | CP | MR | HR | S | HP | CF |
| Computer | 1.17 ± 0.54 | 3.70, 148.02 | 122.40 | < 0.001 | 0.754 | | -1.14 | -1.09 | -1.01 | -1.53 | -6.46 |
| Mechanoid | 2.21 ± 1.24 | | | | | *** | | 0.33 | 0.09 | -0.51 | -2.62 |
| Humanoid | 1.81 ± 0.94 | | | | | *** | ns | | -0.22 | -0.86 | -3.89 |
| Stranger | 2.07 ± 1.22 | | | | | *** | ns | ns | | -0.59 | -2.78 |
| Human | 3.21 ± 1.54 | | | | | *** | ** | *** | *** | | -1.42 |
| Close Friend | 5.67 ± 1.05 | | | | | *** | *** | *** | *** | *** | |

Note: CP = Computer Partner, MR = Mechanoid Robot, HR = Humanoid Robot, S = Stranger, HP = Human Partner, CF = Close Friend. Total N = 42. All p-values reported are Bonferroni corrected, *p<0.05, **p<0.01, ***p<0.001. [§]Greenhouse-Geisser correction.

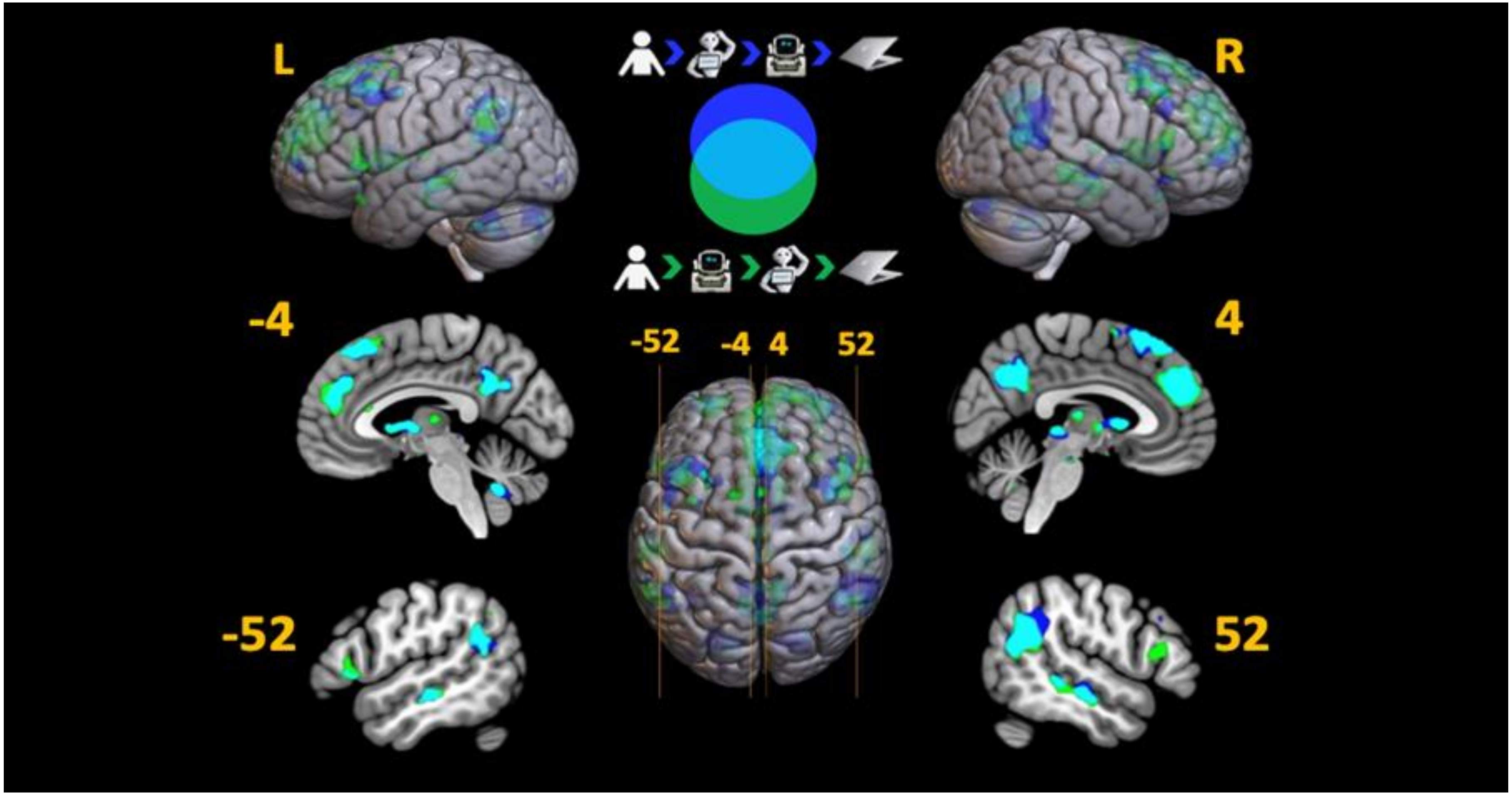


Figure S1. Whole brain T-map overlap analysis Humanness Model (Human > Humanoid > Mechanoid > Computer ; Blue) and the Socialness model (Human > Mechanoid > Humanoid > Computer; Green) - overlap is Teal. p>0.001 unc, no minimal cluster threshold.

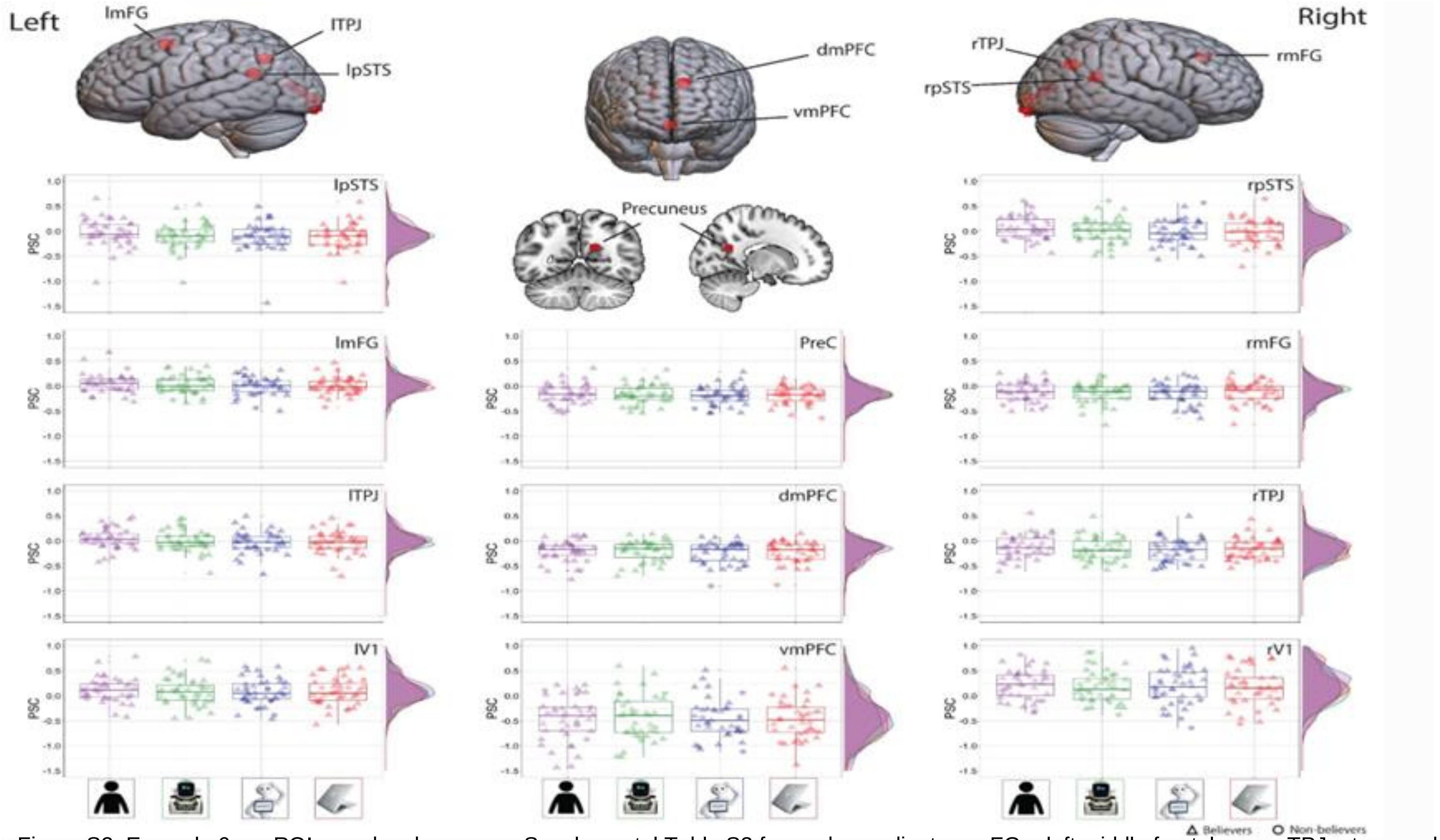


Figure S2. Example 6mm ROI search spheres; see Supplemental Table S3 for peak coordinates. mFG = left middle frontal gyrus; TPJ = temporal parietal junction; pSTS = left superior temporal sulcus; vmPFC = ventromedial prefrontal cortex; dmPFC = dorsomedial prefrontal cortex; PreC = precuneus. Figures represent the distribution of percent signal change (PSC) across the 4 game partners in each ROI. Boxplot show 1st-3rd quartiles. Individual data points with triangles represent “true believers”; those data points with a circle represent “non-believers.”

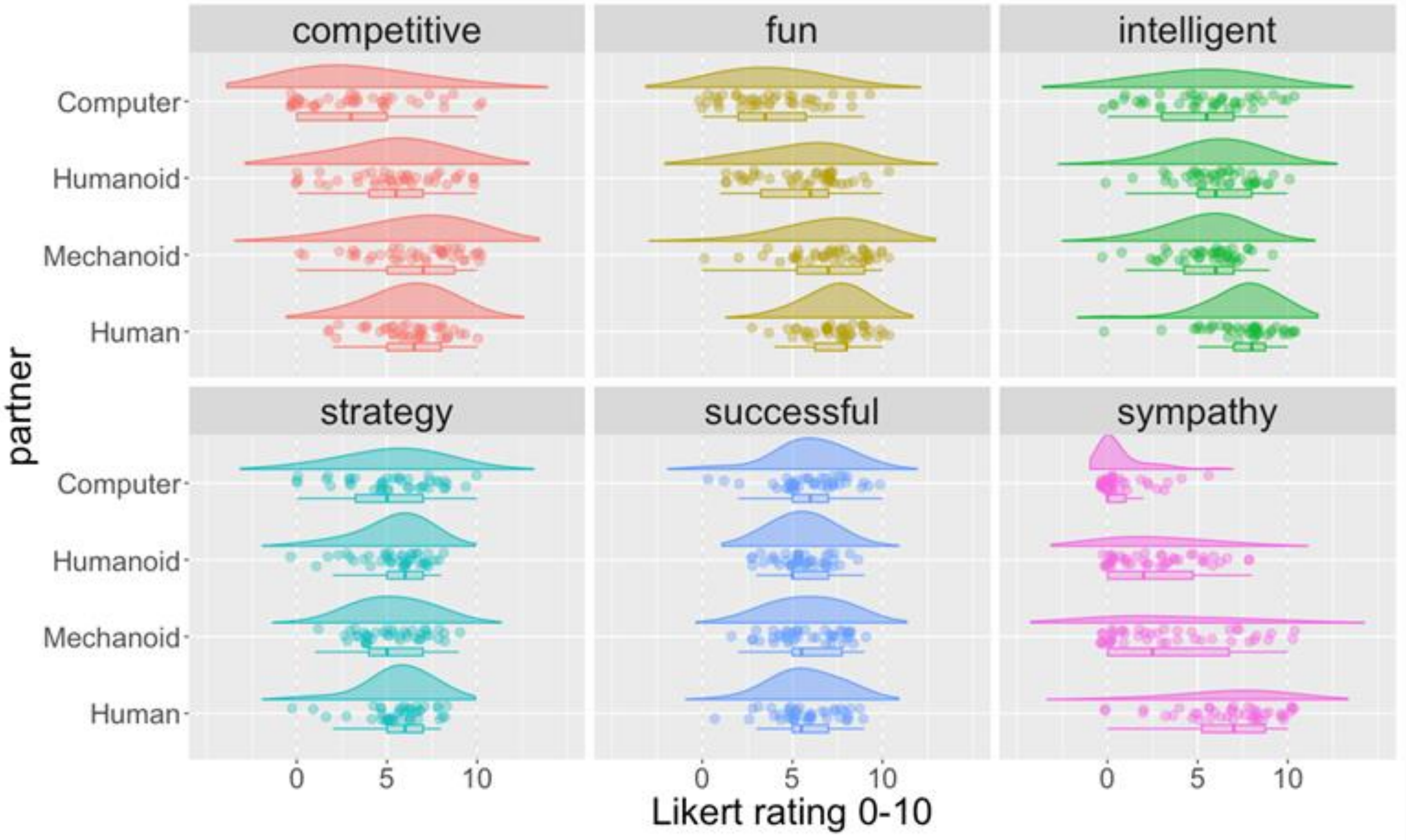


Figure S3. Raincloud plots showing the distribution of rating responses for each of the six debrief questions. Boxplot show 1st-3rd quartiles.
